# GlycoDraw: A Python Implementation for Generating High-Quality Glycan Figures

**DOI:** 10.1101/2023.05.20.541563

**Authors:** Jon Lundstrøm, James Urban, Luc Thomès, Daniel Bojar

**Affiliations:** Department of Chemistry and Molecular Biology, University of Gothenburg, Gothenburg, Sweden. Wallenberg Centre for Molecular and Translational Medicine, University of Gothenburg, Gothenburg, Sweden

**Keywords:** glycan, glycobiology, Python, SNFG, visualization

## Abstract

Glycans are essential to all scales of biology, with their intricate structures being crucial for their biological functions. The structural complexity of glycans is communicated through simplified and unified visual representations according to the Symbol Nomenclature for Glycans (SNFG) guidelines adopted by the community. Here, we introduce GlycoDraw, a Python-native implementation for high-throughput generation of high-quality, SNFG-compliant glycan figures with flexible display options. GlycoDraw is released as part of our glycan analysis ecosystem, glycowork, facilitating integration into existing workflows by enabling fully automated annotation of glycan-related figures and thus assisting the analysis of e.g., differential abundance data or glycomics mass spectra.

## Introduction

Glycans play a crucial role in the biology of all organisms. Structural complexity arises from the presence of a wide range of monosaccharides that can exist in different anomeric configurations and be connected through various linkages. Further, unlike linear biological sequences such as DNA or protein, glycans usually contain branched structures. (Varki 2017).

When choosing a nomenclature for representing glycan structures, researchers must compromise between achieving chemical accuracy while ensuring ease of interpretation, balancing the trade-offs and advantages of each approach. As chemical compounds, all glycans can be described by the International Union of Pure and Applied Chemistry (IUPAC) notation (McNaught 1997). However, this notation proves impractical for daily use, due to the verbosity of the language. Other commonly used glycan nomenclatures include GlycoCT (Herget et al. 2008) and Web3 Unique Representation of Carbohydrate Structures (WURCS) (Tanaka et al. 2014) that both were developed with the aim of optimized machine readability in mind. In contrast, IUPAC-condensed, a simplified version of the standard IUPAC notation, offers excellent human readability and remains widely used for easy communication of glycan structures (McNaught 1997). Further, glycans in the IUPAC-condensed nomenclature can even directly be utilized in computational workflows with glycowork (Thomès et al. 2021), where they are converted into graphs in the back-end, or processed on-the-fly into SMILES strings using software such as GlyLES (Joeres et al. 2023), for applications where exact chemical information is beneficial.

Still, for larger and more complicated glycan structures, the IUPAC-condensed notation is not immediately trivial to interpret, particularly for non-experts. To alleviate this issue, the glycobiology community has decided on standardized guidelines for graphical representation of glycan structures to promote efficient communication. The Symbol Nomenclature for Glycans (SNFG) represents commonly occurring monosaccharides via a combination of colors and shapes, including rules for their assembly into glycan structures (Neelamegham et al. 2019; Varki et al. 2015). A comparative overview between the different mentioned glycan nomenclatures is shown in Table 1.

**Table 1.**
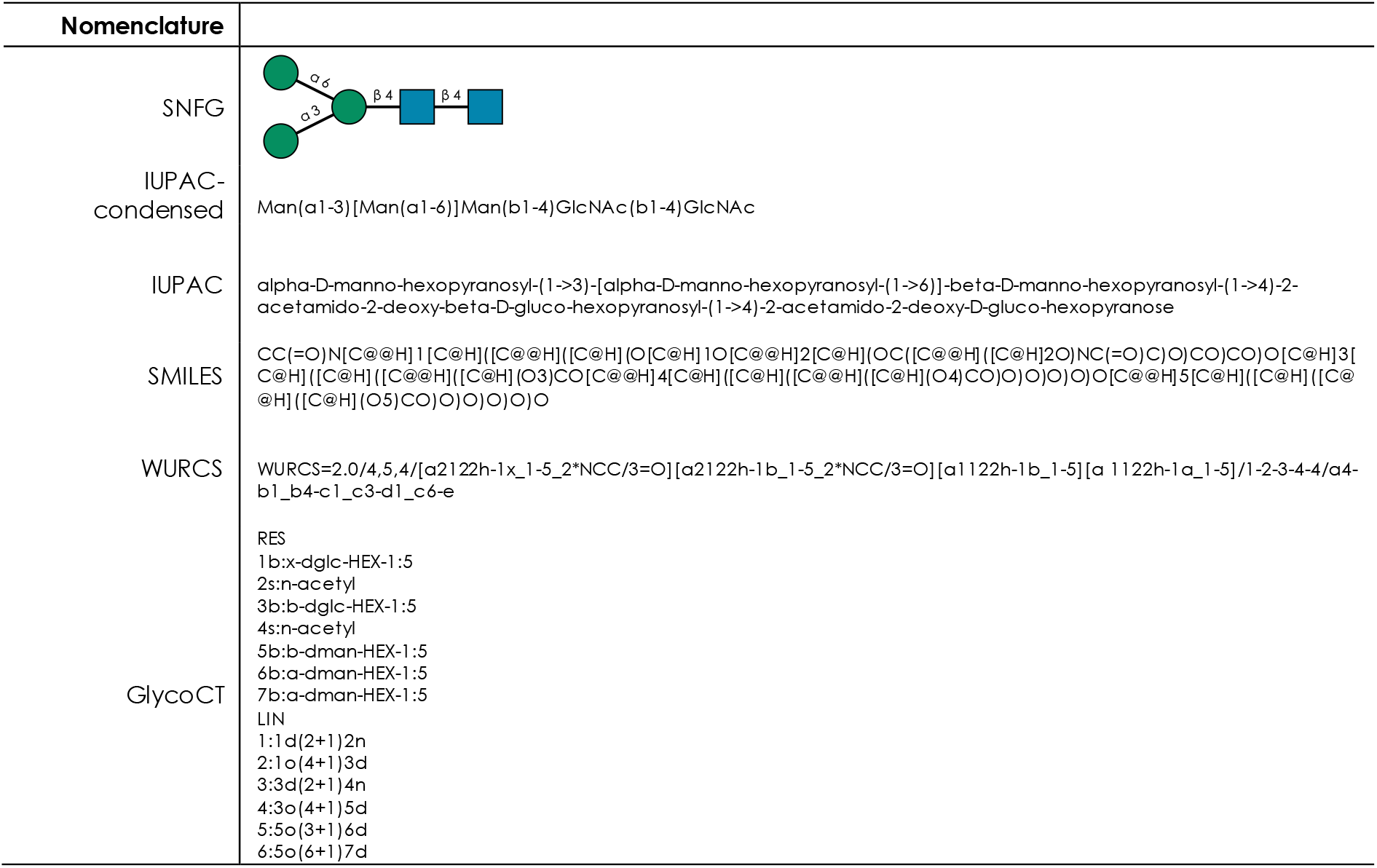
Comparison of glycan nomenclatures.

In recent years, several tools facilitating the generation of glycan figures have been published (exhaustively reviewed in (Lal et al. 2020) and summarized in Table 2). A major focus of available software has been the development of easy-to-use tools with beginner-friendly graphical user interfaces (GUIs). Such GUI-based ‘click-and-drag’ implementations facilitate easy construction of complex structures but are limited with regards to throughput, due to the requirement for manual input. Alternatively, tools such as DrawGlycan-SNFG (Cheng et al. 2017, 2020) or GlycoGlyph (Mehta & Cummings 2020) allow for the generation of glycan figures directly from text (e.g., IUPAC-formatted glycans). However, web-based implementations still limit automatability and scalability. Programmatical implementations exist, e.g., the Python integration of DrawGlycan-SNFG or glypy (Klein & Zaia 2019), but workflow integration possibilities remain limited. In summary, we conclude that there are substantial benefits to filling the current need for a scalable, programmatic, high-quality implementation that can be used in a plug’n’play manner to augment existing workflows.

**Table 2.**
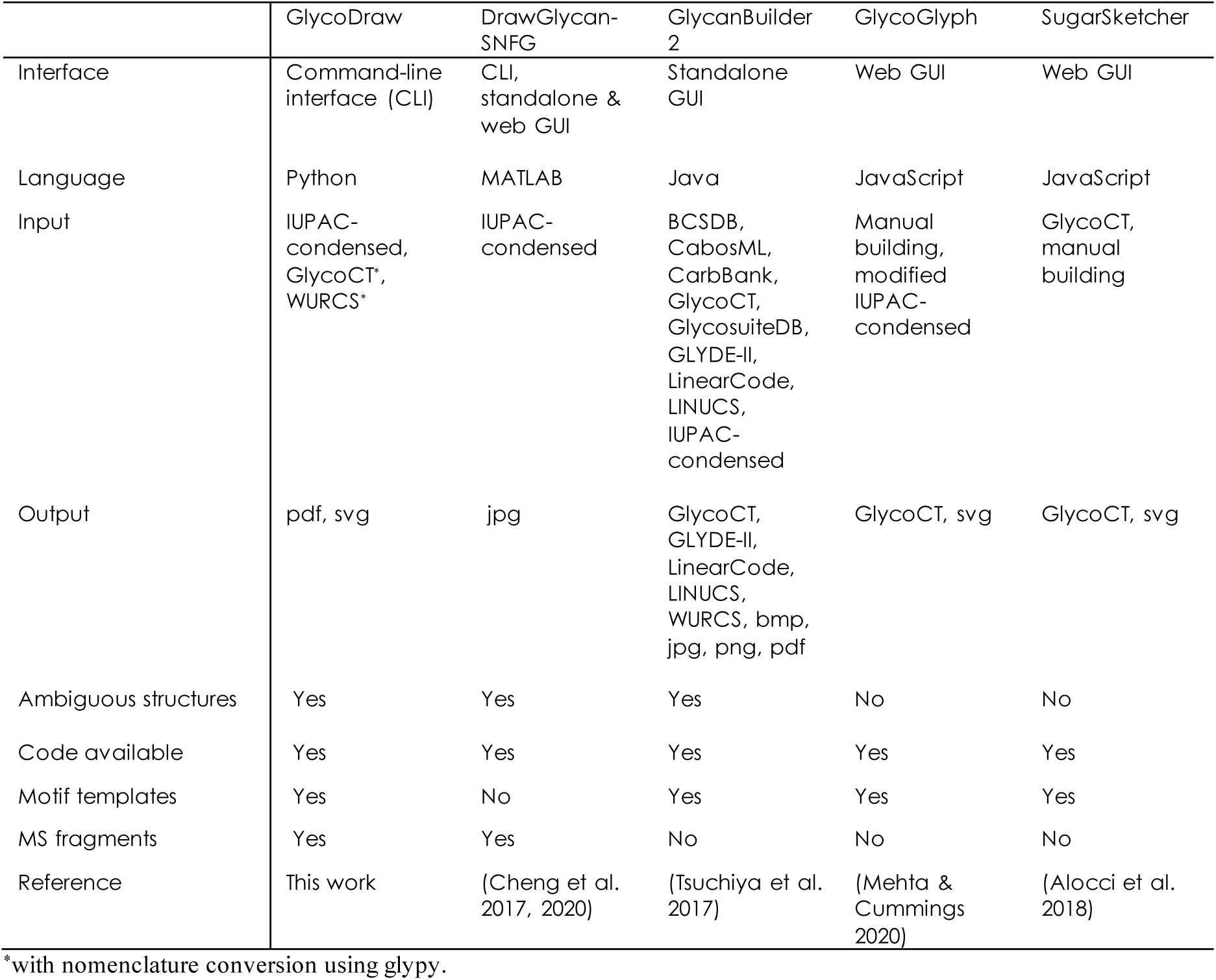
Feature summary of GlycoDraw and other commonly used glycan visualization tools

To overcome these limitations and realize the potential for this computational aspect of glycobiology, we set out to develop a Python-native implementation for high-throughput generation of publication-ready glycan figures, as vector graphics, according to the SNFG guidelines. Here, we describe GlycoDraw, its use in the automation of drawing glycan figures, and integration into existing workflows. GlycoDraw is available as part of glycowork version 0.7, a Python-based ecosystem for glycan analysis we developed and maintain.

## Results

The functionality of GlycoDraw is based on the computational framework of glycowork and the graphics engines of pycairo (version 1.23.0) & drawsvg (version 2.1.1). Glycan inputs in the IUPAC-condensed nomenclature (see Supplementary Note 1 for best practices) are converted to graph objects for calculating coordinates to place individual monosaccharides in relation to each other. The full glycan structure is split into the main chain (i.e., longest chain), branches, and nested branches, while connectivity information is retained based on the glycan graph object. In re-assembling the full glycan as a figure, each individual component is first placed horizontally at the correct x position, as described below, with subsequent adjustment to achieve the proper y position (Fig. 1A).

**Figure 1.**
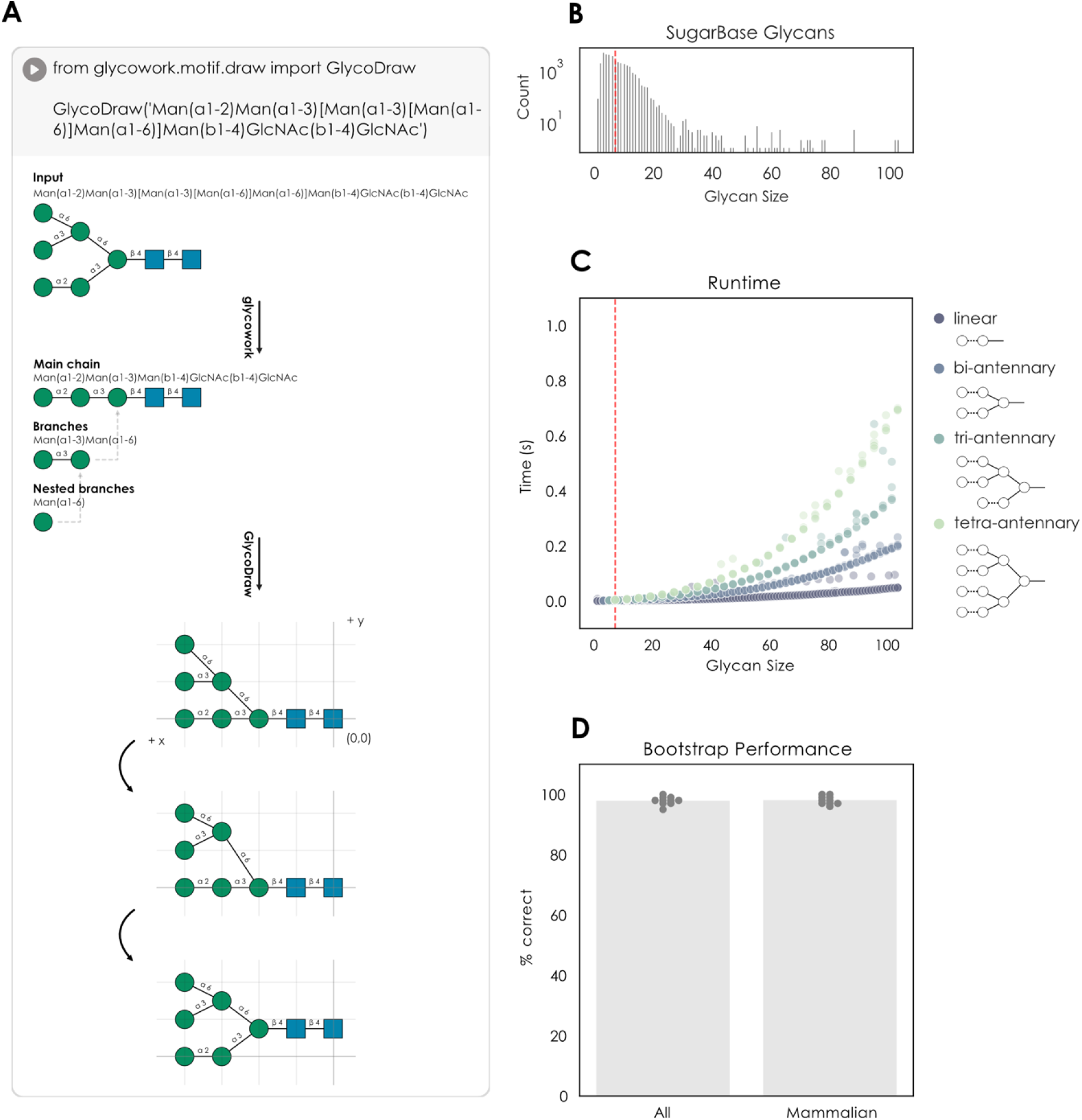
GlycoDraw algorithm and characterization. **(A)** Overview of the drawing algorithm. A glycan string in IUPAC-condensed nomenclature is first split between main chain, branches, and nested branches. Then, monosaccharide coordinates are calculated, and the final figure is rendered. **(B)** Size distribution of glycans in the SugarBase database. The red dotted line indicates the average glycan size (in number of monosaccharides) **(C)** Runtime of GlycoDraw with different glycan topologies and sizes. An average glycan (red dotted line) is rendered in ∽ 2 ms. **(D)** Bootstrap performance of GlycoDraw. 100 glycans from SugarBase (either the full database or limited to mammalian glycans) were selected at random, visualized with GlycoDraw, and the results manually evaluated. This was repeated 10 times for each group of glycan structures.

For the x position, the reducing end (initiating) monosaccharide of the main chain is placed at (0,0) in a coordinate system extending up and to the left. The remaining monosaccharides of the main chain are placed with an incrementally increasing x position (x+1, 0). For branches, the horizontal positioning is calculated in a similar manner, with the coordinates of the initiating monosaccharide being x+1, where x is the position of the monosaccharide that the branch is connected to. For each level of branching, the y position is increased by 1. Next, the y positions are fine-tuned in three steps: (i) any overlap caused by perpendicular monosaccharides (e.g., fucose or xylose) is fixed, (ii) at each branch point, the hub monosaccharide is placed centrally between the branch connections, and (iii) extra space between branches introduced by step (ii) is minimized where possible, to have no more than one unit of distance between branches (Fig. 1A). Finally, the glycan figure is rendered with drawsvg, where monosaccharide icons and chemical modifications are shown according to the guidelines described by the SNFG (described below). GlycoDraw supports inline viewing of the resulting figure when running in a Jupyter Notebook environment, in addition to directly saving figures as pdf or svg files.

In contrast to existing software, GlycoDraw is ideally suited for high-throughput and automatic generation of glycan figures, or situations where the user wishes to integrate figure generation into a computational workflow. While the drawing algorithm scales non-linearly with glycan size and -complexity, the runtime for an average glycan remains around 2 ms, allowing for near-instantaneous rendering of hundreds of structures (Fig. 1B-C). In testing GlycoDraw on the entirety of SugarBase, the glycowork-internal database of nearly 50,000 glycans, we achieved a drawing coverage of 99.5 %, only failing in case of the presence of non-standard monosaccharides not defined in the SNFG. We further tested the performance of GlycoDraw by selecting 100 random glycans from SugarBase and manually evaluating the results for correct formatting. Averaged over 10 iterations (i.e., 1,000 glycans in total), the structure is drawn correctly in 97.9% of cases for all glycans and 98.1% of cases for mammalian glycans (Fig. 1D), with the rare occurrence of minor aesthetical deviations in particularly exotic or complex structures.

As GlycoDraw is implemented within glycowork, it requires Python 3.8+. The GlycoDraw input format is IUPAC-condensed, and the software can handle all canonical SNFG-defined monosaccharides (Fig. 2A). For broad applicability, we also included the *canonicalize_iupac* function (glycowork.motif.processing; version 0.7), which is able to streamline many irregularities or non-canonical formatting of IUPAC-condensed glycans. In addition to the standard glycan figure nomenclature as defined by the SNFG, GlycoDraw supports rendering of glycan fragment ions following the Domon-Costello nomenclature as described in (Domon & Costello 1988) and implemented in GlycoWorkBench (Ceroni et al. 2008) (Fig. 2A). In combination with *in silico* fragmentation, this allows for automated annotation of experimental mass spectra as described below. Monosaccharide modifications are shown according to SNFG guidelines; a furanose ring configuration is indicated by an italicized *f* in the center of the monosaccharide symbol. Likewise, the enantiomeric configuration (D or L) can be shown in the same manner. Chemical carbohydrate modifications are indicated above the monosaccharide symbol, in text, by the number of the modified carbon (or, if unknown, the atom linking the modification), followed by the abbreviated chemical substituent (Fig. 2A).

**Figure 2.**
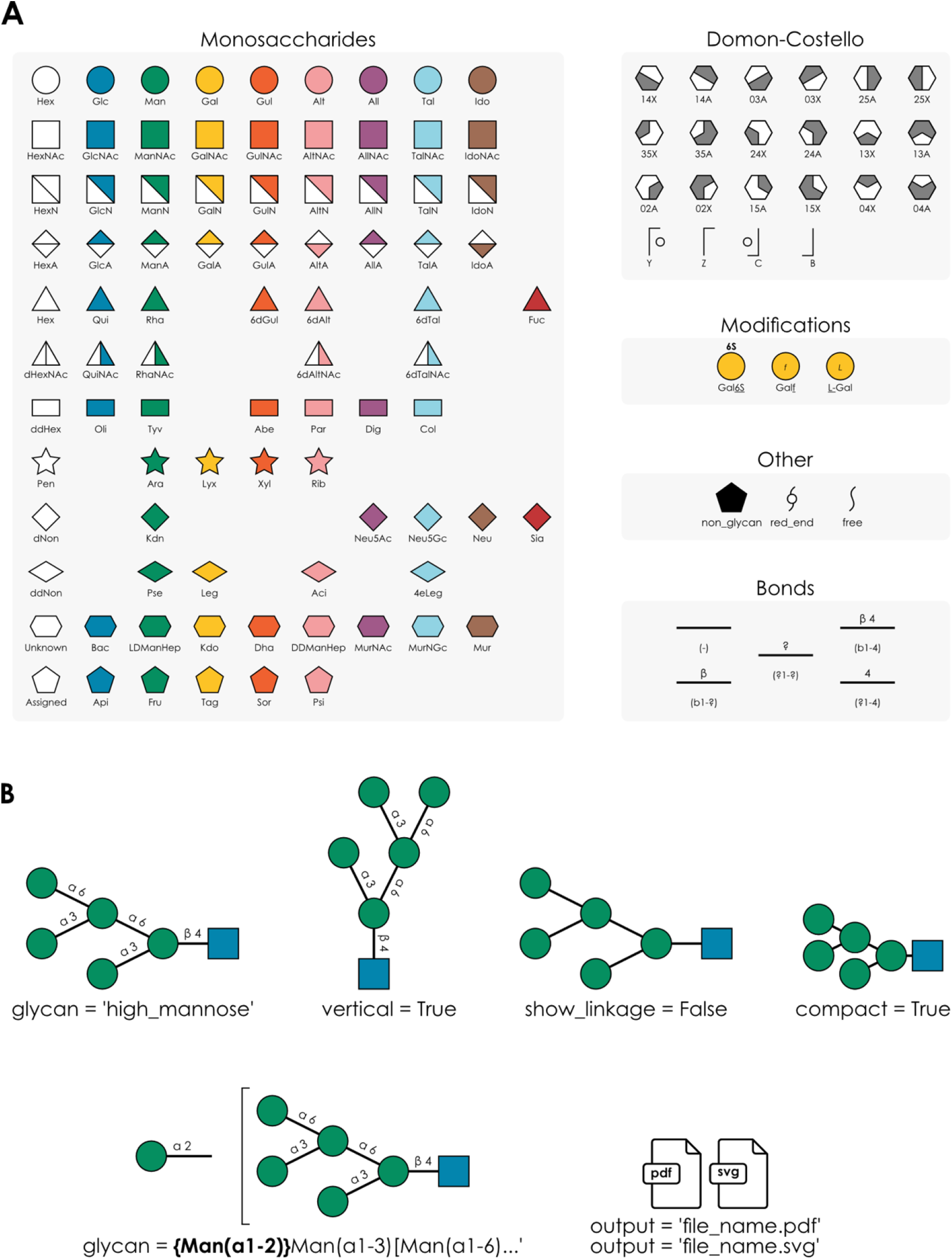
GlycoDraw functionality. **(A)** Overview of SNFG monosaccharide and Domon-Costello fragment icons supported by GlycoDraw. Modifications are shown with the example of galactose and extend to any monosaccharide. Bonds are shown with the example of beta-linkage. **(B)** Optional arguments to GlycoDraw allow modification of the output as well as saving the figure as an image file.

GlycoDraw supports the drawing of a list of predefined glycan structures and -motifs by name (e.g., “LewisX”; *motif_list*, imported from glycowork.glycan_data.loader, contains the full list of possible structures) (Fig. 2B). Further, several optional arguments are available in order to modify the resulting glycan figure. ‘vertical = True’ will draw the glycan is a portrait format rather than the default horizontal format. ‘show_linkage = False’ will hide all linkage information and display a skeletonized version of the glycan backbone, while ‘compact = True’ additionally minimizes the bond length to show the structure as compact as possible (Fig. 2B). Ambiguous glycan structures, e.g., when fucose is present but the exact location in the glycan is unknown, or sialylation of one of multiple branches in a complex *N*-glycan, are also supported by GlycoDraw. The ambiguous part of the structure is enclosed within curly brackets ‘{Fuc(a1-3)}’ and written before the glycan. Best practices dictate that the ordering of multiple ambiguous components follow that of branch ordering, i.e., longest > lowest linkage > alphabetical (Fig. 2B). The output of GlycoDraw can be viewed inline when working in a notebook format, e.g., Google Colaboratory or Jupyter Notebooks. Image files can be saved by specifying “output = ‘filename.pdf’”. Files can be exported as pdf or svg by indicating the appropriate file extension in the name (Fig. 2B), and can be freely edited as vector graphics in downstream applications. A quick-start guide describing the installation and general use of GlycoDraw can be found online in the documentation of glycowork (https://bojarlab.github.io/glycowork/).

The Python implementation of glycan figure generation allows for easy integration with existing glycan analysis and processing workflows. With GlycoDraw, any list of glycans can be effortlessly converted into figures and saved as high-quality vector graphics. Furthermore, automated workflows can be established, in which glycan-related svg figures generated using the Python matplotlib library are modified to include glycan graphics instead of text labels by using the *annotate_figure* function (glycowork.motif.draw; version 0.7) (Fig. 3A). The integration of GlycoDraw with example workflows is described below.

**Figure 3.**
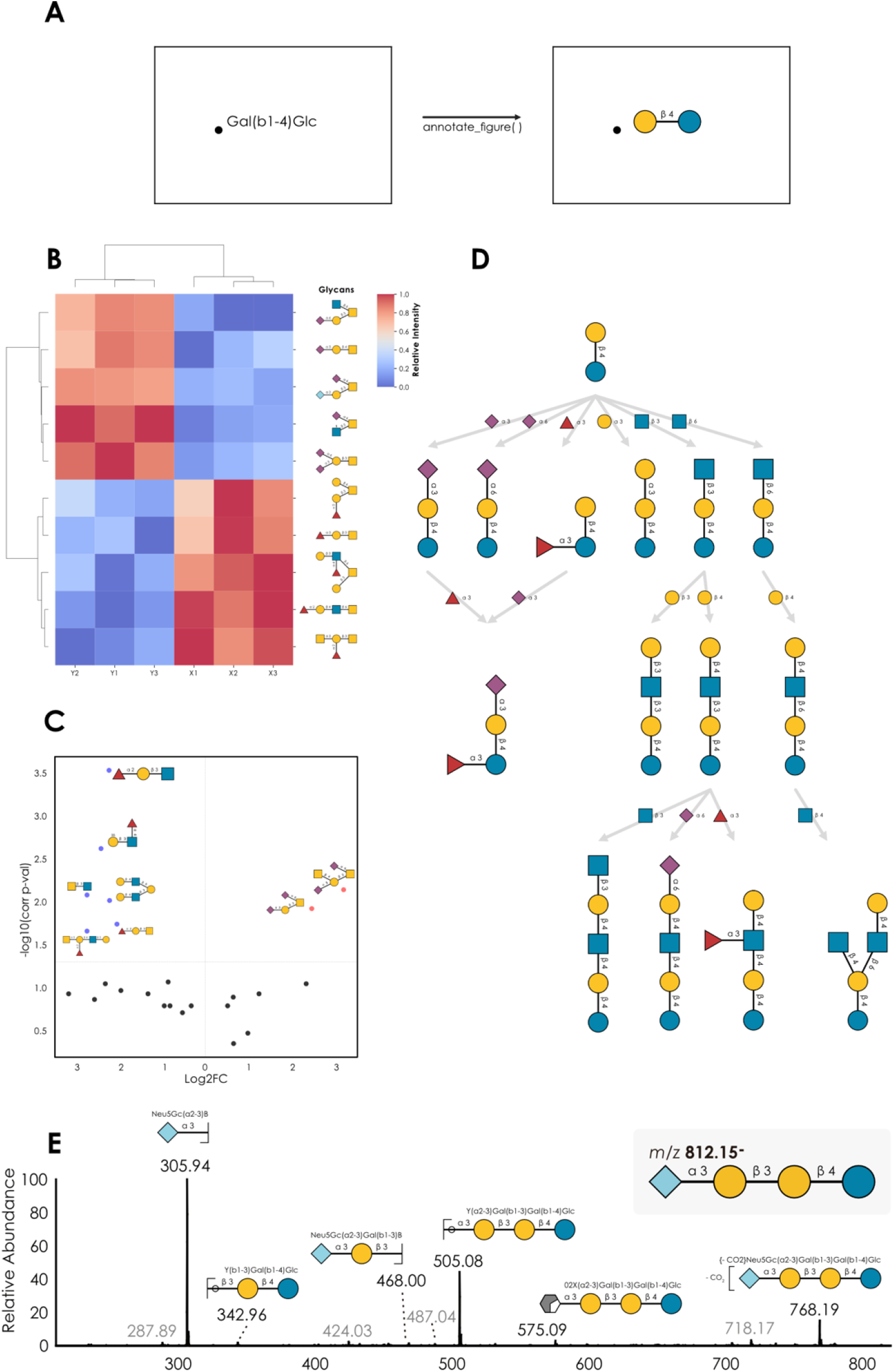
Example applications of GlycoDraw. **(A)** The *annotate_figure* function modifies matplotlib-generated svg figures by replacing text labels with glycan images **(B)** Annotated heatmap visualization of glycan abundance between two groups based on simulated example data. **(C)** Annotated volcano plot visualization of differential glycan abundance quantification based on simulated example data. **(D)** Example milk oligosaccharide biosynthetic network showing the relation between enzymatic reactions and selected structures. **(E)** MS/MS spectrum of Neu5Gc(a2-3)Gal(b1-3)Gal(b1-4)Glc from (Jin et al. 2023), showing the integration of the Domon-Costello nomenclature with GlycoDraw. Low-abundance peaks in grey were not assigned any fragment structure.

Glycowork contains functions to analyze glycan abundance and visualize the results with heatmaps. With the version 0.7 update introducing GlycoDraw, we additionally included functionality to modify such heatmaps by automatic annotation with glycan figures instead of text labels (Fig. 3B). As an alternative to heatmaps, differential glycan (motif) abundance can be visualized with volcano plots, as widely used in the field of RNA-seq and differential expression analysis. Our implementation additionally allows for annotation of volcano plots by adding glycan figures to selected data points, scaled by significance, resulting in easy visual interpretation (Fig. 3C).

Another example of the benefit of automatable glycan figure generation includes the rendering of biosynthetic networks, as described in (Thomès et al. 2023). With glycowork, theoretical biosynthetic networks including all possible intermediate structures can be generated based on the observed glycan abundance of an analyzed sample. Using GlycoDraw, such networks can be annotated to include glycan figures. In the example graph (Fig. 3D), each substructure is shown as the network nodes, and the enzymatic reaction is indicated on the edge connecting two intermediates.

The analysis of glycomics mass spectrometry data routinely includes expert assignment and manual annotation of spectra and fragment mass peaks. For this purpose, glycan fragments are usually annotated using the Domon-Costello nomenclature, a text-based system that can be visualized by specific icons that represent possible fragmentation ions. In addition to standard IUPAC-condensed, GlycoDraw supports this extended fragmentation nomenclature. An example workflow would include computationally obtaining all possible fragments of a specific glycan, cross-checking their masses against a list of observed mass peaks from a glycomics experiment, and generating the corresponding figures where glycan text labels are replaced by glycan images for automatable annotation of glycomics mass spectra (Fig. 3E).

## Discussion

With the introduction of GlycoDraw, we present a Python-native implementation for scalable, high-throughput generation of high-quality, SNFG-compliant glycan figures. GlycoDraw has several advantages over existing tools for generating glycan figures, including scalability for high-throughput generation of images, and importantly, our Python implementation additionally allows for integration into existing workflows, such as the annotation of glycan-related figures.

While we use IUPAC-condensed inputs as the preferred glycan nomenclature in our workflows, we recognize the widespread usage of WURCS and GlycoCT in the community. As a workaround, we refer users to already existing functionality to convert between glycan nomenclatures online (glycosmos.org/glycans/converters, Tsuchiya et al. 2019) or in Python using glypy (Klein & Zaia 2019). As mentioned previously, within different IUPAC “dialects”, such as the formatting of glycan sequences in the CFG array database (www.functionalglycomics.org/static/consortium/CFGnomenclature.pdf), the restructured *canonicalize_iupac* function within glycowork version 0.7 can be used to harmonize inputs. Further, while GlycoDraw functions as expected for the vast majority of glycan structures, we note the existence of very rare cases, particularly exotic/highly complex structures, where manual intervention may be necessary. Fortunately, in such cases the glycan images can be exported as vector graphics and easily adjusted with appropriate image editing software.

GlycoDraw is a valuable tool for generating publication-ready glycan figures that comply with SNFG guidelines. Its scalability and programmability make it ideal for high-throughput analysis. We will continue to develop GlycoDraw to improve its capabilities and usability, and we envision that it will contribute to advancing the field of glycobiology.

## Supplementary Data

Supplementary data are available at *Glycobiology* online.

## Author Contributions

Conceptualization: J.L., D.B., Funding Acquisition: D.B., Resources: D.B., Software: J.L., J.U., L.T., D.B., Supervision: D.B., Visualization: J.L., J.U., L.T., D.B., Writing—Original Draft Preparation: J.L., D.B., Writing—Review & Editing: J.L., J.U., L.T., D.B.

## Funding

This work was funded by a Branco Weiss Fellowship – Society in Science awarded to D.B., by the Knut and Alice Wallenberg Foundation, and the University of Gothenburg, Sweden.

## Conflict of Interest Statement

The authors declare no competing interests.

## Data Availability Statement

All used code and data can be found at https://github.com/BojarLab/glycowork/, with particular emphasis on the motif.draw module.

## Notes

https://bojarlab.github.io/glycowork/

### Competing Interest Statement

The authors have declared no competing interest.

### Summary of Updates

Table 2 updated

