## Supplemental Note for "GlycoDraw: A Python Implementation for Generating High-Quality Glycan Figures"

### Supplementary Note 1 - Best Practices for Writing Glycans in the IUPAC-condensed Nomenclature

Briefly, the IUPAC-condensed format for representing glycans involves showing the sequence of monosaccharides connected by glycosidic bonds. Linkages are denoted in parentheses immediately following a monosaccharide, indicating the position of the anomeric carbon (alpha or beta) of the preceding monosaccharide, followed by the numbered carbons of the preceding and subsequent monosaccharides connected via the glycosidic bond. If there is uncertainty about any elements of the linkage, a "?" is used to replace the uncertain position. Monosaccharides are noted with their common name, and the enantiomer (D or L) is only specified as a prefix if it is the uncommon version. The pyranose form is assumed by default (for all monosaccharides except apiose), while the furanose form is specified with an 'f' suffix. Branches are indicated by placing brackets before the monosaccharide that the branch is connected to. The longest contiguous glycan chain is the main chain, and shorter side chains are branches. If branches are not the same length, longer branches come first. If branches are of equal length, the main chain contains the branch with the lower connection number in its final linkage (e.g., a1-3 > a1-6). If multiple branches are connected to the same monosaccharide, branches of the same length are ordered first by the connection number in their final linkage and then, if needed, alphabetically by the final non-reducing end monosaccharide of the branch. Monosaccharide modifications are indicated by the number of the modified carbon, followed by the modification, and multiple modifications are arranged in ascending order by carbon number. We note that small to moderate deviations from these best practices, particularly regarding branch order, usually do not cause errors in the described glycowork functions.
